# Dehydrated snakes reduce postprandial thermophily

**DOI:** 10.1101/2023.04.05.535800

**Authors:** Jill L. Azzolini, Travis B. Roderick, Dale F. DeNardo

## Abstract

Transient thermophily in ectothermic animals is a common response during substantive physiological events. For example, ectotherms often elevate body temperature after ingesting a meal. In particular, the increase in metabolism during the postprandial period of pythons - known as specific dynamic action – is supported by a concurrent increase in preferred temperature. The objective of this study was to determine whether hydration state influenced digestion-related behavioral thermophily. Sixteen (8 male and 8 female) Children’s pythons (*Antaresia childreni*) with surgically implanted temperature data loggers were housed individually and provided a thermal gradient of 25-45 °C. Body temperature was recorded hourly beginning 6 days prior to feeding and for 18 days post-feeding, thus covering pre-feeding, postprandial, and post-absorptive stages. Each snake underwent this 24-day trial twice, once when hydrated and once when dehydrated. Our results revealed a significant interaction between temperature preference, digestive stage, and hydration state. Under both hydrated and dehydrated conditions, snakes similarly increased their body temperature shortly after consuming a meal, but during the later period of the postprandial stage, snakes selected significantly lower (~1.5°C) body temperature when they were dehydrated compared to when they were hydrated. Our results demonstrate a significant effect of hydration state on postprandial thermophily, but the impact of this dehydration-induced temperature reduction on digestive physiology (e.g., passage time, energy assimilation) is unknown and warrants further study.

**Summary statement:** Dehydration suppresses the extent to which python increase body temperature after ingesting a meal, thus demonstrating a physiological conflict between optimizing body temperature and water balance.

## Introduction

One of the most widely supported tenets of thermal biology is that at all levels of biological organization performances are influenced by temperature, and that performances are executed optimally at a specific temperature or narrow temperature range (T_o_; Brattstrom, 1965; Huey and Bennett, 1987; Block, 1994). Thus, there is an incentive for animals to regulate their body temperature (T_b_) so that they can perform optimally in their environment and, in doing so, increase their likelihood of survival and reproduction (Huey and Hertz, 1984; Huey and Bennett, 1987). Since the optimal temperature for different performances can differ, the target temperature often fluctuates based on current physiological demands (Brett, 1971; Claireaux et al., 1995; Blouin-Demers and Weatherhead, 2001). Optimal temperature for critical physiological processes can be higher than it is during standard metabolism when the organism is supporting embryonic development (Wallman and Bennett, 2006; Lourdais et al., 2008; Lorioux et al., 2012) and digestion (Wiiters and Sievert, 2001; Wang et al., 2002; Tattersall et al., 2004; Wall and Shine, 2008; Raviv and Gefen, 2021) or lower to support sperm production (Cejko et al., 2016) and torpor (Gaertner et al., 1973; McAllan and Geiser, 2014).

While there are clear performance benefits of careful regulation at an optimal temperature, conflicts can influence the extent to which T_o_ and T_b_ are matched. Thermal heterogeneity and other environmental limitations are known to impact heat exchange and thus thermal optimization of a performance (Brett, 1971; Blouin-Demers and Weatherhead, 2001). Additionally, thermophilic behavior to accommodate physiological activities that have higher optimal temperatures may be impacted by limited capabilities or tradeoffs with other physiological needs (Stephens and Krebs, 1986; Secor and Diamond, 2000; Sunday et al., 2014). For example, given that body temperature and evaporative water loss are highly correlated (Mautz, 1982; Guillon et al., 2014), reduced water availability may create a tradeoff between thermal optimization of physiological performance and limiting water loss. Despite the well-established relationship between water loss and body temperature, most studies of ectotherm thermoregulation have not tested for the presence of synchronous thermo-hydroregulation processes (Rozen-Rechels et al., 2019). Accordingly, limited water availability may play a large role in an animal’s dynamic thermophilic response to optimize a particular performance.

Digestion is a vital physiological process as it enables an organism to obtain energy and other important nutrients (Lignot et al., 2005; Ott and Secor, 2007; Cox et al., 2008). During digestion, ectothermic animals have an elevated metabolism (Secor and Diamond, 1998), which is referred to as “specific dynamic action” (SDA, Ott and Secor, 2007), and concomitantly, there is often a postprandial elevation in T_b_ (Secor and Phillips, 1997). This postprandial elevation in T_b_ likely results from a combination of metabolic heat production associated with the increase in metabolism and altered behavioral thermoregulation (Witters and Sievert, 2001; Wang et al., 2002; Wall and Shine, 2008; Raviv and Gefen, 2021). Since an animal will lose more water as T_b_ increases, postprandial thermophily may exaggerate the physiological conflict between energy balance and water balance, especially if water availability is limited. As the majority of terrestrial environments inhabited by organisms experience some degree of water limitation, at least at a seasonal scale (Hao et al., 2018), it is prudent to understand the tradeoff between energy and water balances, and to what extent, if any, does water imbalance influence the thermal response to feeding. While there is evidence that some insectivores and herbivores gain a hydric benefit from eating (Cooper, 1985; Degen et al., 1997, Ostrowski et al., 2002), the opposite has been shown for binge-feeding reptiles. Despite the considerable water content of large whole-animal meals, consuming such meals does not benefit (Wright et al., 2013) and may even worsen (Murphy and DeNardo, 2019) dehydration in binge-feeding squamates. Consistent with this finding, snakes provided with ready access to water drink more after consuming a meal (Lillywhite, 2017), further indicating a potential hydric cost of digestion.

Children’s pythons (*Antaresia children;* Gray, 1845) serve as an excellent study system for understanding the extent to which hydration state can influence thermophilic responses to optimize specific physiological performance. This species naturally experiences extended annual dry seasons that can lead to water imbalance (Brusch et al., 2017), and thus faces the need to perform while in a dehydrated state. Additionally, Children’s pythons are binge feeders that only intermittently consume relatively large meals, providing the opportunity to easily evaluate individuals under distinct postprandial and post-absorptive conditions.

This study addressed whether hydration state influences the dynamics of thermal adjustments made during digestion. We hypothesized that dehydration suppresses a behavioral thermophilic response during digestion. We predicted that pythons would have the highest T_b_ during the first 96 hours following a meal, the timeframe during which postprandial metabolic rate peaks (Fig. 1). Further, we predicted the increase in T_b_ would slowly decrease and return to baseline as digestion approaches completion. Finally, we predicted that throughout digestion, pythons would prefer higher temperatures when in a hydrated state compared to when they are dehydrated.

**Figure 1.**
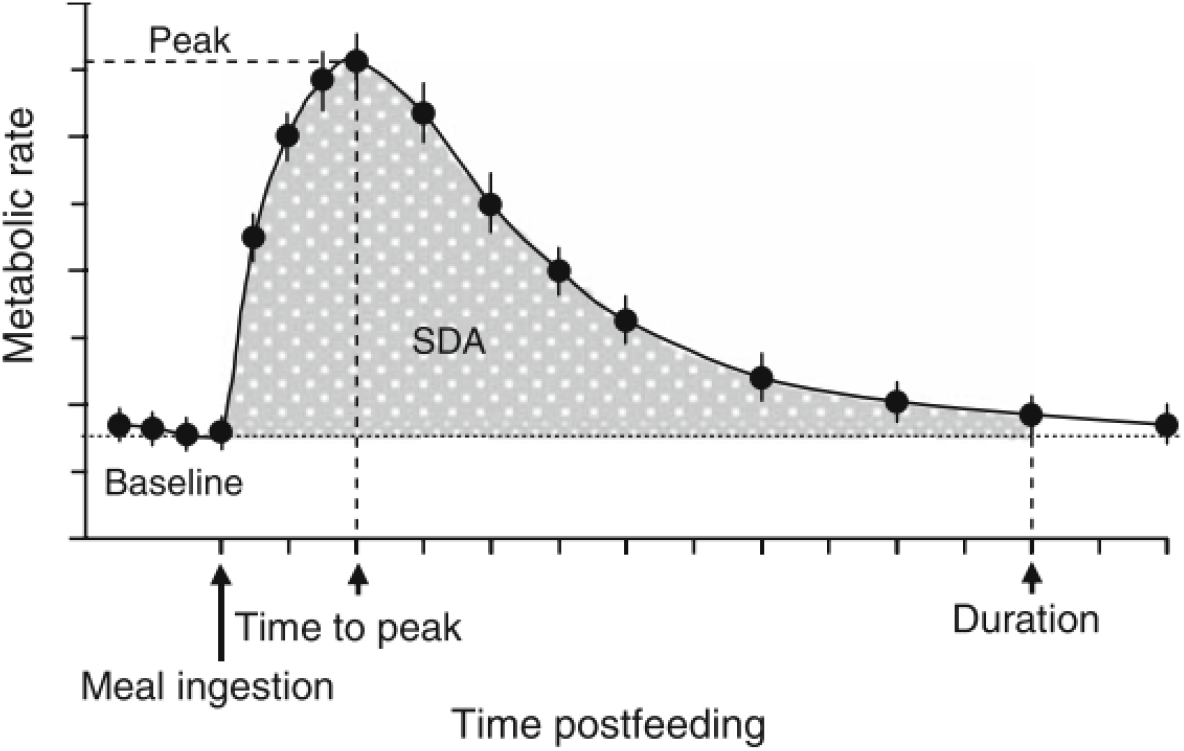
Visual representation of the rise in metabolic rate following meal ingestion, known as Specific Dynamic Action (SDA), modified from Ott and Secor, 2007.

## Materials and methods

### Study Organism

The Children’s python is a medium-sized (up to 1.2 m snout-to-vent length, SVL, 600g mass) constrictor native to the wet-dry tropics of northern Australia (Wilson and Swann, 2003) where they experience natural annual fluctuations in water resource availability. Free-standing water can be locally absent for 3-4 months at a time, typically between May and August (Taylor and Tulloch, 1985). All Children’s pythons used in the study were captive-bred individuals that have been maintained as part of a long-standing colony at Arizona State University. All experimental procedures were approved by the Arizona State University Institutional Animal Care and Use Committee (protocol 20-1740R).

### Experimental Design

The experiment was designed to examine the effect of dehydration on T_b_ when the snakes were post-absorptive (i.e., not digesting a meal) and postprandial (i.e., during meal digestion). Sixteen adult Children’s pythons (n = 8 males; average mass 506 ± 3 g; range 494-520 g; average SVL 95.9 ± 1.5 mm; range 90.6-102.4 mm; n = 8 females; 550 ± 5 g; 531-571 g; 95.7 ± 1.6 mm; 89.3-100.9 mm) were housed individually in semilucent plastic drawer cages that had tops made of expanded metal (30 cm D x 18 cm W x 10 cm H, Freedom Breeder, Turlock, CA). Room temperature was set at 25 ± 0.5°C and subsurface heating (Flexwatt, Flexwatt Corp., Wareham, MA) was provided at the rear of each cage so that the cage provided a 25-45°C thermal gradient. A piece of semi-rigid, highly absorbent paper (Techboard, Shepherd Specialty Papers, Watertown, TN) covered the bottom of the cage so that the snake could choose to be exposed on the surface or secluded under the paper regardless of the temperature selected. The snakes were randomly distributed among six rows of a rack, three cages per row, with an additional row of cages above and below these six rows to ensure that, for consistency, all snakes had a cage row above and below them. Prior to the start of the experiment, all snakes were provided water ad libitum but were not fed once placed on the experimental rack.

Snakes were given a week to acclimate to their cages, and then temperature loggers (Thermochron iButtons #DS 1922L, Maxim Integrated Products, San Jose, CA) programmed to record temperature (± 0.05°C) hourly and coated with Plasti-Dip (PDI Inc, Woodcliff Lake, NJ) were implanted intracoelomically following the methods of Lourdais et al., 2008. While under isoflurane anesthesia, a logger was secured to the body wall just caudal to the gall bladder using non-absorbable suture (Braunamid, B. Braun Medical, Melsungen, Germany) to ensure the logger remained stationary.

Snakes were given two weeks to recover from their surgeries before they were weighed using a platform scale. To balance the size and sex of snakes between two groups, snakes of each sex were ordered from heaviest to lightest and then alternatingly assigned to two groups. That is, the heaviest male was assigned to Group 1 (G1), the second and third heaviest to Group 2 (G2), the fourth and fifth heaviest to G1, etc. This was repeated for females, so that there were 4 males and 4 females in each group. The two groups were differentiated by the order in which they would experience the two hydration states (i.e., G1 snakes experienced the hydrated condition first, whereas G2 snakes experienced the dehydrated condition first). Once assigned to their groups, each snake was randomly assigned to a location on the housing rack.

Dehydration was accomplished by withholding food and water for 30 days. This duration was chosen as it causes a moderate level of dehydration and would enable the ensuing 24-day feeding cycle to be completed in a total of 54 days without water, which approximates the duration of water deprivation for Children’s pythons in a previous study (Dupoué et al., 2014). As G2 underwent the dehydrated conditions first, they were given 6 days of water access between ending the dehydrated cycle and beginning the hydrated cycle. Throughout the 54-day dehydration period, snakes were weighed weekly to ensure body mass did not drop more than 15%. During the hydrated cycle, water bowls were checked daily to ensure water was always available.

### Body temperature assessment

Once all snakes had completed testing under both the hydrated and dehydrated conditions, the temperature loggers were removed following the anesthesia and surgery protocol described for the implantation surgery except that, upon entering the coelomic cavity, the anchoring suture was cut and the logger removed from the snake. The hourly temperature data were downloaded from the loggers and then analyzed to compare the average T_b_ of multiple digestive stages during both hydrated and dehydrated states (Fig. 2). We assigned five stages of the feeding cycle for use in our analyses. “Pre-feeding” data was defined as the average hourly body temperature for the 6 days immediately preceding the day of feeding. “Postprandial” data were collected for 12 days after the snake was fed a 30 ± 1 g thawed mouse (5.9% body mass of male snakes, 5.5% body mass of female snakes). Our rationale for the allotted 12 days was that metabolism in pythons, as in many snakes, rises relatively rapidly after consumption of a meal and then tapers off more slowly over subsequent days, reaching pre-feeding levels approximately 12 days post-feeding (Ott and Secor, 2007). We divided the 12-day period into three 4-day periods: postprandial days 1-4, postprandial days 5-8, and postprandial days 9-12. This separation of postprandial timepoints incorporated peak digestive effort (days 1-4 post-feeding), a time when metabolism can be ten-fold more than pre-feeding levels (Ott and Secor, 2007; Fig. 1), as well as the gradual decline from peak effort to baseline. Following the completion of the postprandial periods, we collected “post-absorptive” data, which consisted of the average hourly T_b_ collected on days 13-18 post-feeding. This 24-day cycle breakdown into five periods was then duplicated for the data collected when the snake was under the alternative hydration state (dehydrated for G1 and hydrated for G2).

**Figure 2.**
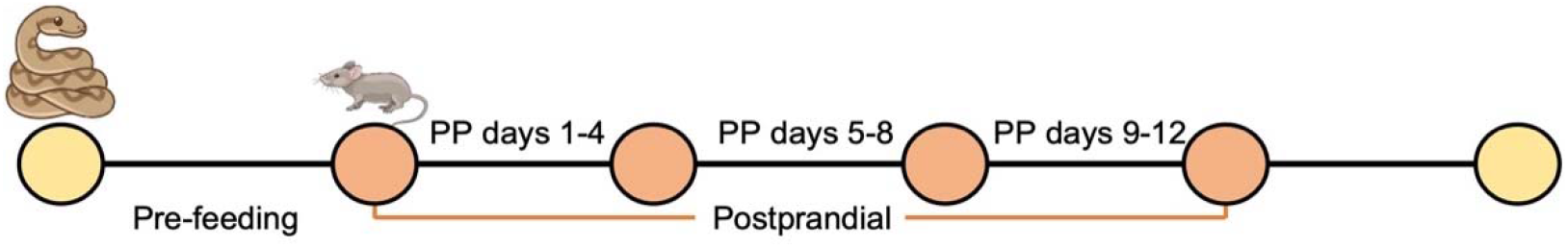
Digestive stages compared in this study. Days 1-6 leading up to a meal represented “pre-feeding” (PF), The “postprandial period” (PP) started after the snake had been fed and continued for 12 days, which were further divided into 4-day intervals (e.g., “PP days 1-4”) to distinguish metabolic difference during the postprandial period. Lastly, the “post-absorptive” (PA) stage covers days 19-25. Snakes completed the timeline twice: once when they were provided water *ad libitum* and the other after being water-restricted for 30 days prior to and then throughout the ensuing timeline.

### Statistical analysis

We performed all statistical analyses using RStudio Version 1.0.153. We chose a linear mixed effects model using the “lme4” package since our data contained continuous covariates and possible nonlinear relationships. Due to our small sample size, we set our statistical significance at 0.10 for all models and post-hoc analyses. We tested for an interaction between average temperature, feeding stage, and hydration status. We included sex and body condition index (BCI; calculated as residuals of a body mass vs. SVL regression) as fixed effects. We also included snake ID as a random effect. We tested for normality using the Shapiro-Wilke test (p = 0.56) and assessed multicollinearity using Variance Inflation Factors (all VIFs < 2.5, “car” package). We used Estimated Marginal Means (aka Least-squares means, “emmeans” package) for post hoc testing of pairwise comparisons between digestive stage and hydration status.

To evaluate intra-individual thermoregulatory precision, we evaluated the standard error of the mean (s.e.m.) of each animal’s b T_b_ over the course of each feeding stage as a dependent variable. The model used for this analysis had the same fixed and random effects as the model used to analyze inter-individual temperature preference. Again, we tested for normality by using the Shapiro-Wilke test (p = 0.87), tested multicollinearity using Variance Inflation Factors (all VIFs < 2.5), and used Estimated Marginal Means for post hoc testing comparisons.

## Results

### Body Mass

Throughout the dehydration process, G2 snakes lost 63 ± 9 g (range 23-103 g), or 13% body mass while G1 snakes lost an average of 58 ± 5 g (range 43-72 g), or 11% body mass. Upon being given water at the end of the dehydration cycle, G2 snakes returned to within 30 ± 11 g (range 4-56 g; 94%) and G1 snakes returned to within 36 ± 2 g (range 29-44 g; 93%) of their pre water deprivation mass within 24 hours of being provided water but no food. This supports previous work indicating that the vast majority of mass lost during water deprivation periods in snakes is due to water loss (Dupoué et al., 2015). Following the completion of the hydrated feeding cycle, G1 snakes were at 98% (12 ± 6 g, range −9-30 g) and G2 snakes within 99% of their initial body mass (7 ± 4g, range −29-5 g).

### Body temperature - within a given hydration state

All 16 snakes completed the hydrated feeding cycle. We found that hydrated snakes had significantly elevated T_b_ during postprandial days 1-4 compared to pre-feeding T_b_ (p = 0.0072, Fig. 3) as expected due to the increased metabolic demands of digestion. T_b_ during postprandial days 5-8 was intermediate in that it was not significantly different from any of the other periods (all p > 0.10). The average T_b_ during postprandial days 1-4 was significantly higher than postprandial days 9-12 (p=0.074) and the post-absorptive period (p = 0.013). Postprandial days 9-12 and post-absorptive average Tb’s were not significantly different from pre-feeding body temperature (p = 0.99, p = 1.000).

**Figure 3.**
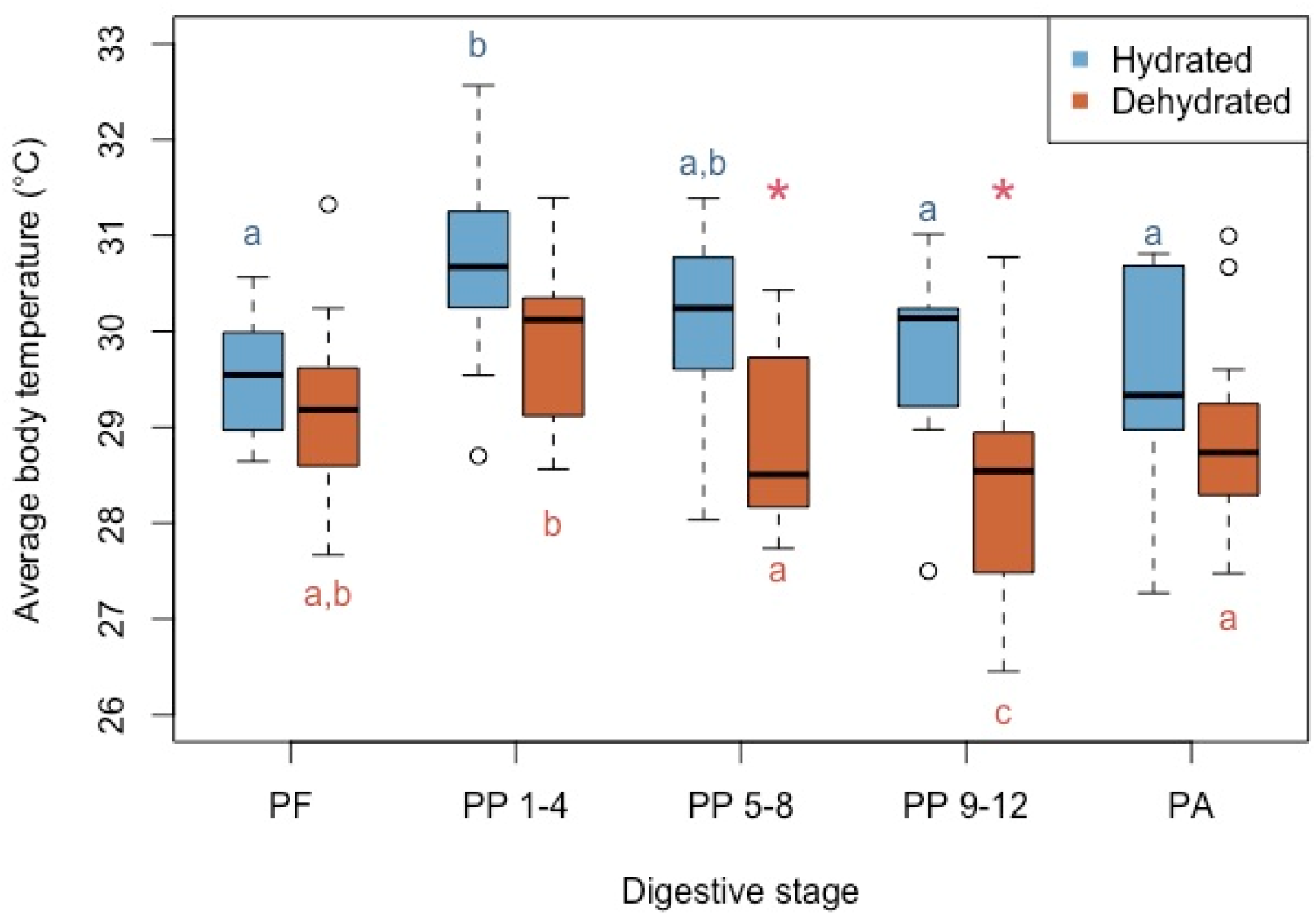
Average T_b_ of all pythons during the various feeding stages (pre-feeding, PF; postprandial, PP; post-absorptive, PA) when the snakes were hydrated (n = 16, blue boxes) and dehydrated (n = 13, orange boxes). The bold line inside each box represents the mean, the top and bottom of each box represent the 75^th^ and 25^th^ quartiles, respectively. Whiskers denote the s.e.m, and open circles denote outliers. The blue letters reflect significant differences among the hydrated measurements, while orange letters show significant differences among the dehydrated measurements. Asterisks denote significant differences between hydrated and dehydrated animals at a given feeding stage.

Only 13 snakes completed the dehydrated feeding cycle. One snake died shortly after the end of the hydrated conditions phase. The cause of death was unknown; however, it did not appear to be linked to the assigned treatment as the snake had only been without water for eight days. We also had two snakes that refused to eat during their dehydrated feeding cycle. All three snakes were in G1, the group that experienced the hydrated feeding cycle before the dehydrated feeding cycle.

Postprandial thermal dynamics during dehydration were similar to those when hydrated (Fig. 3). Dehydrated snakes chose significantly warmer T_b_ during postprandial days 1-4 than they did during postprandial 5-8 (p = 0.086), postprandial 9-12 (p < 0.0001), and post-absorptive (p = 0.047) periods (Figure 3). Similarly, post-absorptive T_b_ was not significantly different from pre-feeding T_b_ (p = 0.99). Unlike when they were hydrated, the average pre-feeding T_b_ when the snakes were dehydrated was not significantly different than T_b_ during the postprandial 1-4 period (p = 0.23).

### Body temperature - between hydration states

There was a significant effect of both hydration status (p < 0.0001) and digestive stage (p < 0.0001) on T_b_. The average T_b_ during postprandial 5-8 and postprandial 9-12 periods were lower when snakes were dehydrated compared to when they were hydrated (−1.1°C, p = 0.027 and −1.5°C, p = 0.0006, respectively; Fig. 3). Thus, snakes reduced their postprandial T_b_ more quickly when dehydrated.

There was also a significant effect of digestive stage (p = 0.0029) and hydration (p = 0.0099) on the precision of intra-individual T_b_ over time. That is, T_b_ showed significantly less within individual variation during the dehydrated pre-feeding measurements than during the dehydrated postprandial days 5-8 (p = 0.042) and at every point during the hydrated postprandial period (days 1-4, p = 0.0009; days 5-8, p = 0.0029; days 9-12, p = 0.017.

### Body temperature - between sexes

We found a significant effect of sex (p = 0.018) on T_b_ that persisted regardless of hydration state. In all stages other than postprandial days 1-4 and whether hydrated or dehydrated, males were, on average, 0.63°C warmer than female snakes (Fig. 4A and 4B).

**Figure 4.**
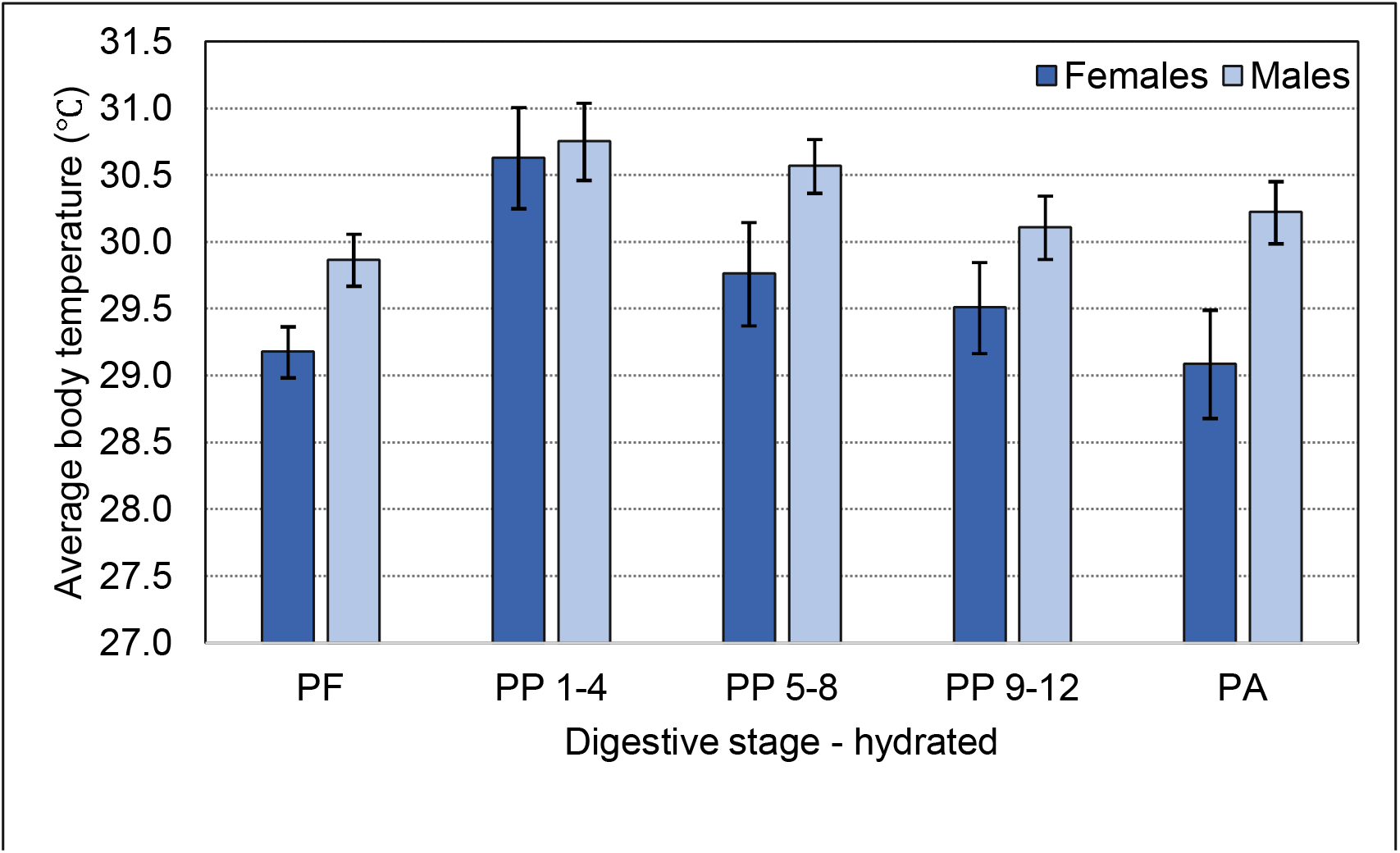
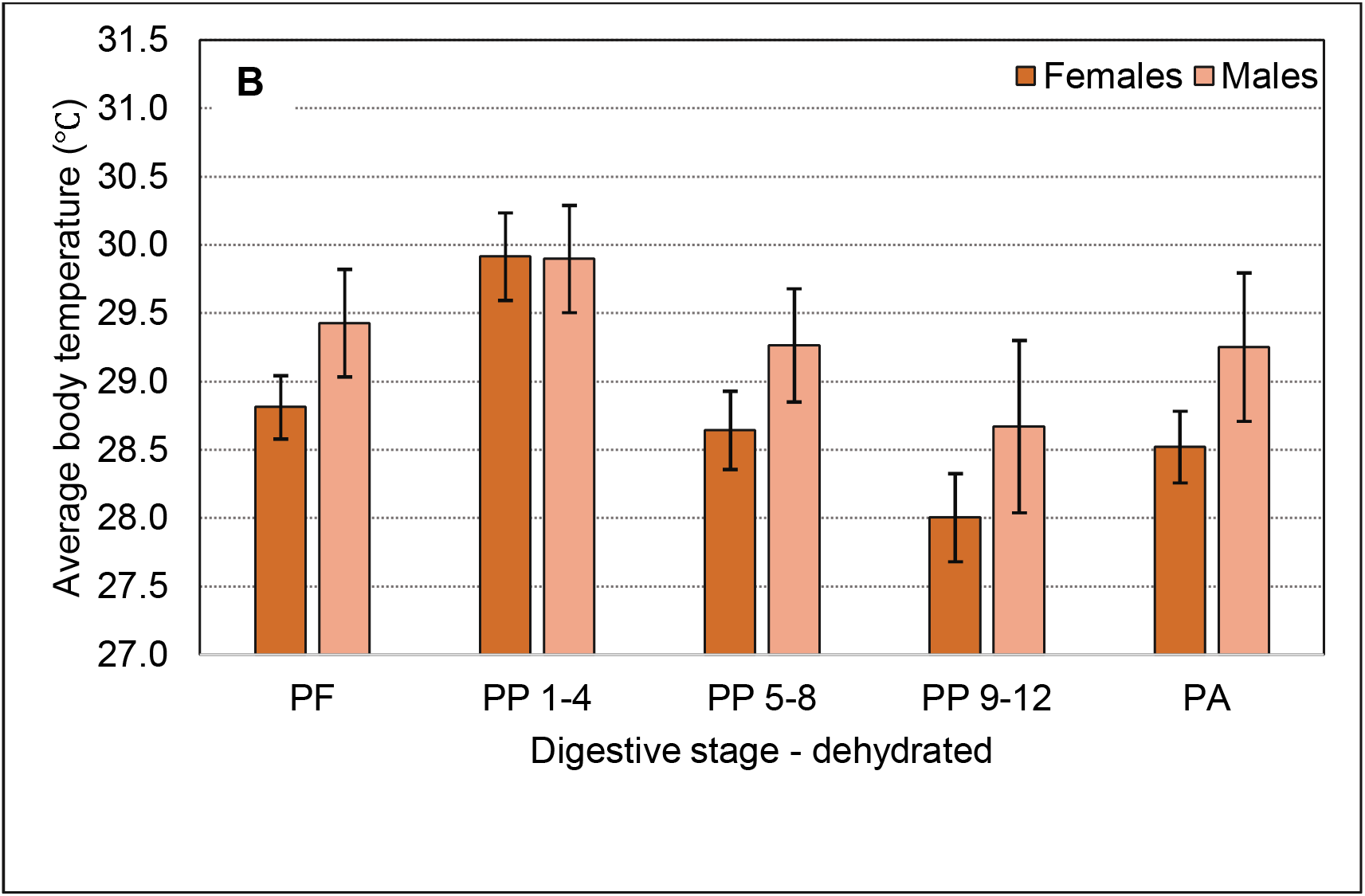
T_b_ differences between the sexes when (A) hydrated and (B) dehydrated throughout the respective feeding cycles. The top of each bar depicts the mean T_b_, and the error bars represent s.e.m.

## Discussion

In both hydrated and dehydrated conditions, python T_b_ peaked during postprandial days 1-4, supporting our prediction that pythons would choose the highest T_b_ during the first 96 hours following a meal regardless of hydration state (Figure 3). Our results complement existing research regarding behavioral thermophily during digestion in ectotherms. This phenomenon has been documented in a variety of different snake species, including rat snakes (*Elaphe obsoleta obsoleta*), carpet pythons (*Morelia spilota*), common water snakes (*Nerodia sipedon*), and common garter snakes (*Thamnophis sirtalis*) (Blouin-Demers and Weatherhead, 2001). In another carnivorous, binge-feeding ectotherm, the Gila monster (*Heloderma suspectum*), postprandial thermophily was positively correlated with meal size (Gienger et al., 2013). Gila monsters not only had higher T_b_ as their meal size increased, they also maintained higher T_b_ for up to twice as long when given a meal that was 20% of their body mass versus a meal that was only 5% of their body mass (Gienger et al., 2013). Beyond squamates, there is substantial diversity in species that become thermophilic during digestion. For example, sculpin (*Cottus extensus;* Wurstbaugh and Neverman, 1988) migrate to warmer water after feeding, and Woodhouse’s toads (*Bufo woodhousiĩ*) increase body temperature following meals whereas fasted toads do not change body temperature over time (Witters and Sievert, 2001). Postprandial thermophily has also been documented in invertebrates such as leeches (*Hirudo verbena;* Petersen et al., 2011) and scorpions (*Hottentotta judaicus;* Raviv and Gefen, 2021).

As predicted, after peaking there followed a progressive decrease in T_b_ through postprandial days 5-8 and 9-12 (Fig. 3). The trend in our temperature data mimics the metabolic curve seen in a typical SDA response (Figure 1), where the peak metabolic rate is reached within four days of meal consumption. Following this peak, the metabolic rate slowly returns to the baseline rate (Secor and Diamond, 1998). In one study that compared metabolic rates between 8 species of snakes, some of which naturally are frequent feeders and others, like Children’s pythons, are infrequent feeders, all snakes reached their peak metabolic rate within 4 days of meal consumption (Secor and Diamond, 2000). Further, infrequently feeding species in this study had the largest factorial increase in metabolic rate, ranging from 10-to 18-fold increases in VO_2_ from fasting to postprandial (Secor and Diamond, 2000). The similar shapes of the body temperature response curve in our study and the classic SDA curves are likely related, at least in part, to endogenous heat production related to digestive activity. While thermogenesis during digestion has not been documented in Children’s pythons, increased metabolic heat production during digestion in the South American rattlesnake, *Crotalus durissus*, increased body temperature by 1.2°C (Tattersall et al., 2004). While increased metabolism likely accounts for some increase in T_b_, behavioral thermophily likely provides a significant contribution to increased T_b_ during digestion. For example, wild rattlesnakes (*C. atrox, C. molosus*, and *C. tigris*) fitted with intracoelomic temperature-sensing radio transmitters were observed retreating to shelter immediately following meal consumption, but within 24-72 hours most snakes were found fully basking (Beck, 1996). The few snakes in this study that were not basking had adjusted position within their shelter so that they were partially exposed to sunlight. Basking snakes had an average T_b_ of 31°C, whereas the unfed snakes’ average T_b_ was only 25°C. Unlike physiological processes related to locomotion, which can have a broad range of optimal temperatures, the T_o_ for energy assimilation tends to require more precise thermoregulation (Angilletta, 2001). In frequently feeding ectotherms such as eastern fence lizards (*Sceloporus undulatus*), the preferred T_b_ of ~33°C is very close to the T_o_ for digestive efficiency (Angilletta, 2001). Further, lizards captured from locations in Utah, South Carolina, and New Jersey were found to have similar T_b_ despite the differing climates, indicating that the lizards must behaviorally thermoregulate in order to maintain the preferred temperature. Both endogenous heat production related to metabolic activity and behavioral thermophily likely contribute to increased postprandial T_b_ in ectotherms, but the relative importance of metabolic heat production in binge-feeding ectotherms warrants further investigation.

Our final prediction that throughout digestion, pythons would prefer higher temperatures when in a hydrated state compared to when they are dehydrated, was mostly supported. When dehydrated, snakes were significantly cooler during postprandial days 5-8 and 9-12 (Fig. 3). It is notable that T_b_ was not significantly different between hydration states during the postprandial days 1-4, when T_b_ was highest. Although our findings were statistically significant, the physiological significance of a 1.5°C reduction in T_b_ requires further investigation. Pythons have approximately 96% digestive efficiency at a range of temperatures from 24°C to 33°C (Bedford and Christian, 2000), so a 1.5°C temperature reduction may seem insignificant from a functional standpoint. However, elevations in T_b_ require an investment of energy (e.g., for endothermy), commitments of time (e.g., for basking), and/or additional predation risks (e.g., during basking efforts), so thermophily, even at a finer scale, must convey benefits. Irrespective of the thermal sensitivity of energy assimilation, lower T_b_ may extend gastrointestinal passage time (Angilletta, 2001; Raviv and Gefen, 2021). Digestion is a vulnerable state for many vertebrate species (Claireaux et al., 1995; Wang et al, 2000), so even a slightly prolonged digestive process could make wild animals more at risk of predation. Additionally, longer passage time may force longer durations between meals and, for ectothermic animals living in seasonal climates, the ability to consume frequent meals in a limited time window is especially crucial (Secor and Phillips, 1997) and more frequent meals have been shown to improve body condition and reproductive success (Tattersall et al., 2004; Taylor et al., 2004).

Previous work on other vital physiological processes, such as reproduction, have shown an extremely sensitive relationship with T_b_. In one study where river lamprey were maintained at 14°C, 10°C, and 7°C, females held at 14°C were first to release their eggs (Cejko et al., 2016). Even within Children’s pythons, a comparative review of existing data reveals a highly sensitive relationship between T_b_ and gravidity duration. At the pythons’ preferred T_b_ during reproduction (31.3°C), pythons had an average gravidity duration of 23.1 days (Lourdais et al., 2008). In contrast, Lorioux et al. (2012) maintained reproductive pythons at a slightly higher constant temperature of 31.5°C and found the average gravidity duration to be slightly shorter at 22.5 days. Lastly, Azzolini et al. (unpublished data) held reproductive pythons at a constant temperature of 31.0°C and found a slightly longer gravidity duration of 24.8 days. Together, these results demonstrate that a difference of only 0.5°C in T_b_ results in a 2-day difference in gravidity duration in this species. Based on this knowledge, the physiological significance of the 1.5°C reduction in T_b_ during digestion caused by dehydration warrants investigation.

Surprisingly, we found a significant effect of sex on T_b_ during most digestive states, where males were consistently warmer than females. Whether hydrated or dehydrated, males were, on average, 0.63°C warmer than females at all stages except postprandial days 1-4 (Fig. 4A and 4B). As reported above, we similarly did not find a significant effect of hydration state on T_b_ during postprandial days 1-4. Given that postprandial days 1-4 are the time of peak metabolic activity during digestion, it may be critical for pythons to maintain peak T_b_ during this period regardless of hydration state or sex. While we did not anticipate a sex-based difference, there is pre-existing evidence that sex can affect T_b_ and digestive physiology. Male and pregnant female Atlantic stingrays (*Dasyatis sabina*) chose significantly warmer temperatures than did non-pregnant females (Wallman and Bennett, 2006). In Children’s pythons, the assimilation efficiency of some nutrients significantly differed between sexes (Stahlschmidt et al., 2011). The authors attributed this to differences in body composition, where females store more energy in the form of fat whereas males prioritize skeletal muscle mass (Stahlschmidt et al., 2011). Perhaps the temperature preference during digestion influences specific nutrient absorption and disposition; however, that is beyond the scope of our study.

Both water balance and digestion are vital performances, and our study demonstrates a likely tradeoff between the two in terms of selected T_b_ during digestion. Energy and water balances may tradeoff in additional ways. Interestingly, the only two instances where a snake refused a meal were when the individual was dehydrated. Digestion in snakes requires greater water intake (Lillywhite, 2017) and consuming a meal in water-deprived western diamond-back rattlesnakes (*C. atrox*) worsens their dehydration (Murphy and DeNardo, 2019). Therefore, food refusal by two water-deprived pythons in our study may reflect the tradeoff between energy and water balances in that the opportunity to obtain energy resources was bypassed to not exacerbate current water imbalance. Lastly, it is important to note that our presentation of the thermal gradient provided a predictive daily thermal landscape. It is unknown how animals would respond in an environment that presents higher levels of complexity (Wall and Shine, 2008; Burggren, 2019; Nancollas and Todgham, 2022). Future studies could incorporate added factors for individuals to consider when choosing postprandial T_b_, such as proximity of water resources, vulnerability at heat sources, and frequency in the availability of potential prey.

## List of abbreviations used

| List of abbreviations used |  |
| --- | --- |
| Specific dynamic action | SDA |
| Body temperature | T<sub>b</sub> |
| Optimal temperature | T<sub>o</sub> |
| Group 1, Group 2 | G1, G2 |
| Pre-feeding | PF |
| Post-prandial | PP |
| Post-absorptive | PA |

## Acknowledgements

We thank Dr. Stephen Pratt and Dr. George Brusch for their assistance with statistical analyses. We also express our sincere gratitude to the DeNardo lab undergraduate research team for their help with data collection and manuscript review.

## Data availability

Data will be made available through Dryad within one month of publication.

